# LAVASET: Latent Variable Stochastic Ensemble of Trees. A novel ensemble method for correlated datasets

**DOI:** 10.1101/2023.10.20.563223

**Authors:** Melpomeni Kasapi, Kexin Xu, Timothy M.D. Ebbels, Declan P. O’Regan, James S. Ware, Joram M. Posma

**Affiliations:** Section of Bioinformatics, Division of Systems Medicine, Department of Metabolism, Digestion, and Reproduction, Faculty of Medicine, Imperial College London, London, W12 0NN, UK; National Heart & Lung Institute, Imperial College London, London, W12 0NN, UK; MRC London Institute of Medical Sciences, Imperial College London, London, W12 0HS, UK; Royal Brompton & Harefield Hospitals, Guy’s and St. Thomas’ NHS Foundation Trust, London, SW3 6NP, UK; Program in Medical & Population Genetics, Broad Institute of MIT & Harvard, Cambridge, MA, US

**Keywords:** random forest, machine learning, correlation, omics data

## Abstract

**Motivation:** Random Forests (RFs) can deal with a large number of variables, achieve reasonable prediction scores, and yield highly interpretable feature importance values. As such, RFs are appropriate models for feature selection and further dimension reduction (DR). However, RFs are often not appropriate for correlated datasets due to their mode of selecting individual features for splitting. Addressing correlation relationships in high dimensional datasets is imperative for reducing the number of variables that are assigned high importance, hence making the DR most efficient. Here, we propose the LAtent VAriable Stochastic Ensemble of Trees (LAVASET) method that derives latent variables based on the distance characteristics of each feature and aims to incorporate the correlation factor in the splitting step.

**Results:** Without compromising on performance in the majority of examples, LAVASET outperforms RF by accurately determining feature importance across all correlated variables and ensuring proper distribution of importance values. LAVASET yields mostly non-inferior prediction accuracies to traditional RFs when tested in simulated and real 1D datasets, as well as more complex and high-dimensional 3D datatypes. Unlike traditional RFs, LAVASET is unaffected by single ‘important’ noisy features (false positives), as it considers the local neighbourhood. LAVASET, therefore, highlights neighbourhoods of features, reflecting real signals that collectively impact the model’s predictive ability.

LAVASET is freely available as a standalone package from https://github.com/melkasapi/LAVASET.

## Introduction

Random Forest classifiers (RFs) are frequently used to analyse biological data for prediction and feature selection tasks. RFs can deal with a large number of variables, achieve reasonable prediction scores, and yield highly interpretable feature importance values (1). As such, they are appropriate models for feature selection and further dimension reduction (DR) for integrated datasets. The premise of the original RF algorithm is to assemble an ensemble of trees that complement each other and increase variability of predictor selection (2). However, each node and subsequent split still only consider one predictor variable, limiting both the predictive ability and correct feature importance assignment in complex biological settings that include correlated features.

Nguyen et al.(2) have recently developed a new information criterion statistic to evaluate the contribution of features to the predictive ability of the model. It comprises of different categories of probabilities that assess the feature’s proximity to the target class and the complexity of the relationship with the given class. In addition, permutation-based feature importance has been extensively studied in RFs. It has been demonstrated that there is some level of bias in the assignment of feature importance when there exists collinearity between features that are both associated with the target outcome (3).

Few techniques have been proposed for enhancing feature importance calculations in datasets with highly correlated variables. The Boruta algorithm (4) uses a ‘shadow’ feature approach where it duplicates the original dataset and shuffles the feature values. Boruta trains a classifier on the enhanced dataset and assigns a value of importance to each of the features. Then, the algorithm performs a number of iterations where it compares the importance of the real features to the shadow (shuffled) features, while recording how many times the original features outperform the shadow ones (hits). A threshold is set and depending on the number of hits, the respective feature is assigned an importance or removed from the matrix of further iterations. Boruta, in reality, combines permutation importance, by shuffling the original features, with recursive feature elimination (5), by iteratively considering and removing features that do not reach a threshold.

These methods are efficient in removing noisy features that might not reflect real signals, especially by eliminating these through iterations. However, they are not sensitive in picking all relevant features when these are collinear. Addressing relationships between collinear features in high dimensional datasets is imperative for reducing the number of features that are assigned high importance and thereby making the DR more efficient. Here, we propose a novel method termed LAtent VAriable Stochastic Ensemble of Trees (LAVASET) that derives latent variables based on the distance characteristics of each feature and thereby incorporates the correlation factor in the splitting step. Hence, it inherently groups correlated features and ensures similar importance assignment for these. Distance characteristics for the features can include the feature’s adjacent points in a 1D spectrum, adjacent features of time-series data, or spatial distance in 3D structures among other examples. In this context, LAVASET addresses a major limitation in the interpretation of feature importance of RFs when the data are collinear, such as is the case for spectroscopic and imaging data.

## Methods

### A. Datasets

We demonstrate the LAVASET algorithm (detailed below) on four different datasets with feature importances calculated as a result of a prediction/classification problem between disease and healthy control groups or simulated groups.

#### A.1. Irritable Bowel Syndrome - faecal metabolomics (1D)

The Maastricht University Irritable Bowel Syndrome (MIBS) cohort includes human faecal water samples analysed with ^1^H Nuclear Magnetic Resonance (NMR) spectroscopy for 267 participants (146 IBS patients; 121 healthy controls (HCs)). Details on demographics, sample collection, and data acquisition can be found in <u>6</u>. Unlike the original publication, we used the full NMR spectrum (digitised to a total of 18,000 features) following the removal of the internal standard and the water region, and baseline correction.

To test LAVASET’s ability in capturing relevant features, we also created simulated groups from the MIBS cohort dataset. The groups were generated by using 2 (uncorrelated) compounds that each have 2 multiplets, with signals spread out across the length of the spectrum. The compounds used are metabolites ethanol, with peaks at 1.17-1.20 and 3.64-3.68 ppm, and uracil with peaks at 5.79-5.81 and 7.53-7.56 ppm.

#### A.2. Diabetes - urinary metabolomics (1D)

Human urinary metabolomics data from individuals with type-2 diabetes mellitus (T2DM), freely available from Metabolights (<u>MT-BLS1</u>), were used as an additional test cohort. Prior work on this dataset has shown higher classification accuracy compared to IBS data. A total of 84 samples were collected, consisting of 12 healthy volunteers with data at 7 time points, and 30 individuals with T2DM with data collected at 1-3 time points (total of 50 spectra). These were analysed by ^1^H-NMR spectroscopy to evaluate the urine profiles between T2DM and HCs. Metabolite identification was performed by PLS-DA models previously, as described by the authors in <u>7</u>. The raw data were downloaded from Metabolights and processed to standardise each spectrum to 18,000 data points. The water and internal standard regions were removed from the spectrum with the remaining points used for the modelling.

#### A.3. Acute myocardial infarction - electrocardiogram (1D)

Electrocardiogram (ECG) data from the Physikalisch-Technische Bundesanstalt (PTB) dataset (8) were downloaded from PhysioNet (9). We extracted ECG data from individuals with acute myocardial infarction (MI) and compared this against no acute MI. We excluded all data without a reason for admission or with an unknown diagnosis. This resulted in 175 control and 346 acute MI ECGs. The individual ECGs were processed to correct signal drifts. An average cardiac cycle was extracted for each individual that was normalised to 750 data points (0.75s). We used the 3 Frank leads (vector cardiogram, VCG) as input to the algorithm and visualise the feature importance in the conventional 12-leads by making use of the (absolute) Kors regression transformation (10) for the 8 independent leads.

#### A.4. Hypertrophic Cardiomyopathy - CMR imaging (3D)

To test LAVASET’s performance on high dimensional, spatial, 3D datasets, we used data meshes derived from cardiac magnetic resonance (CMR) imaging of the human left ventricle. Segmentation of these images (to produce the meshes) was performed using a deep learning framework developed by collaborators (11). Measurements of myocardial wall thickness were calculated along radial segments that connected the inner endocardial and outer epicardial surfaces (12). This process produced a 46,808 *×* 4 matrix, where for each of the 46,808 points there is a value for the x, y, and z coordinates of the ventricle, and the wall thickness at that point. These segmentations were performed on images in the UK Biobank dataset (13) and an in-house Hypertrophic Cardiomyopathy (HCM) cohort (Royal Brompton Hospital Cardiovascular Biobank (14)). The total number of samples used here was 1,273, consisting of 634 hypertrophic cardiomyopathy (HCM) patients and 639 demographically-matched HCs.

### B. Approach

LAVASET operates given a number of prerequisites and hyperparameters that can be optimised (Figure 1). A user-specified distance matrix is calculated to select the k closest feature points to the feature of interest (FOI), which form the FOI neighbourhood. This FOI neighbourhood submatrix is defined as:

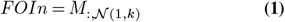

where *M* denotes the matrix containing all the input features and the set 𝒩(1, *k*) represents the indices of the *k* features in *M* that are closest to FOI (based on the user-specified distance matrix), including the index for FOI itself. The user specifies the maximum number of features to consider for each split (with default the square root of the total number of features), and these are randomly selected from the entire dataset in each step. For each selected FOI, the first left singular vector (PC1) of the respective FOI neighbourhood is calculated. We calculate this via Singular Value Decomposition (SVD) of the sub-matrix *FOIn*:

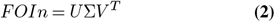

Here, *FOIn* is the scaled FOI neighbourhood for the selected VOI, *U* is the matrix of left singular vectors, Σ is a diagonal matrix containing singular values, and *V* ^*T*^ (or *V* transposed) is the matrix of right singular vectors. The PC1 or first left singular vector is the first vector in matrix *U*, and this is now the latent variable for the FOIn. Loadings for each latent variable are calculated as:

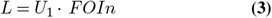

where the first left singular vector *U*_1_ is multiplied by the original *FOIn* sub-matrix. The input matrix for determining the best split will now consist of the latent variable values instead of the original feature values. The best-split variable and value are evaluated by the traditional Gini index method by deducting the sum of the squared probabilities of each class from one. Once the split occurs, the Gini gain is calculated for the selected latent variable and node by subtracting the sum of the Gini index weights of the two child nodes from the parent node. This is repeated recursively until all samples are split into pure leaf nodes, similar to the classic CART algorithm (15).

**Fig. 1.**
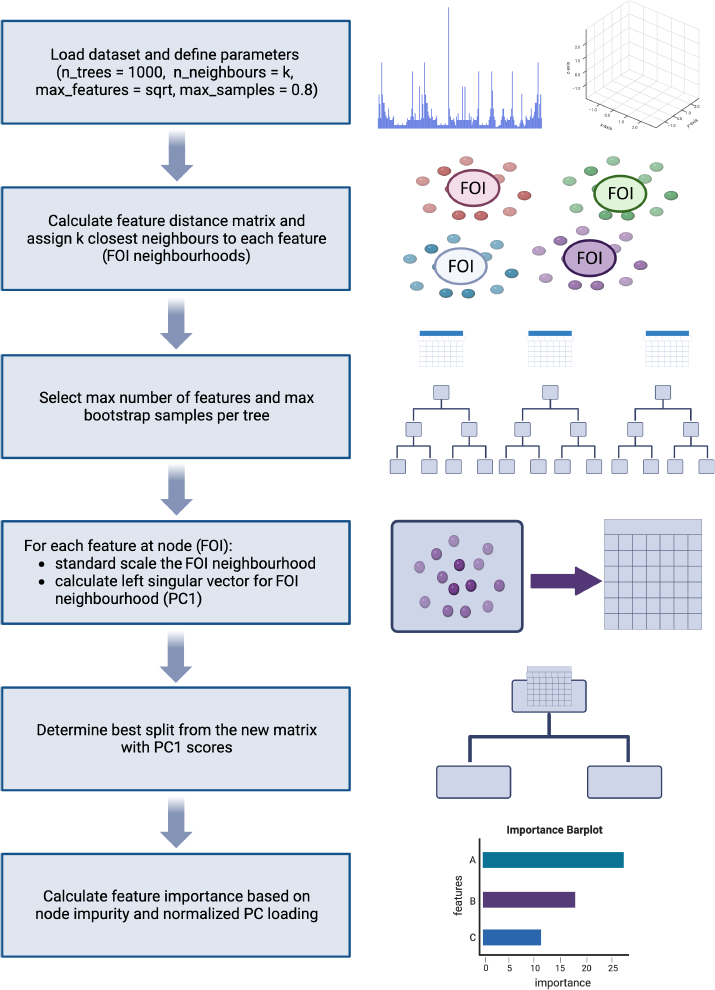
LAVASET high-level pipeline indicating the different and novel approach in the node splitting step and feature importance calculation. Model input is customisable and LAVASET can perform on different types of omics data, from 1D to 3D.

### C. Feature importance calculation

Feature importance scores are calculated for each feature by weighing its contribution to the PC score (left singular vector). Specifically, for every selected feature where the PC score is calculated, the loadings vector of the score (with a shape equaling the number of neighbours considered) is calculated as shown in Equation <u>3</u>. The loadings vector *L* is then multiplied by the Gini gain value assigned to the selected latent variable. This results in a feature importance score not only for the FOI but also for its neighbours.

The LAVASET algorithm is designed to allow for multiple variations in the calculation of a feature’s importance after model construction. LAVASET outputs for each feature_*i*_ a vector of values shown in Equation 4, where 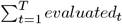 represents the summation over all trees from 1 to *T* of the count of times a feature is evaluated in each tree. Similarly, 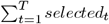 and 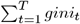 represent the total counts of times a feature is selected for a split and the accumulated Gini values in each tree, respectively. All features of the original matrix *M* are equally *evaluated* for a split with a frequency that follows a Gaussian distribution. The subset of features *selected* for splitting a node are those assigned a feature importance.

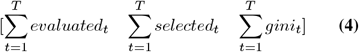

From the values in Equation 4 we can calculate a normalised Gini feature importance by dividing by the sum of all importances. We also evaluate the ratio of the number of times a feature is selected over the count of times it is considered for a split, and incorporate this in our feature importance final value. Using the same output, another use case would be to compare the ratio of Gini values over the count of times a feature is selected for split, which can give an idea of the value magnitude assigned to a feature at a given split. Results presented in this paper utilise the normalised feature importance multiplied by the ratio of 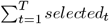 over the 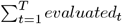 for each feature. This is shown in equation 2 where *importance*(i) is the importance of the feature i, T is the total number of trees, *importance*(i, t) represents the Gini importance of the feature *i* in the tree *t*, and *F* is the total number of features. The sum in the numerator of the first fraction goes over all the trees from 1 to *T* for the feature *i*. The sum in the denominator of the first fraction goes over all the trees and all the features (*f* represents each feature in the feature set), which normalises the Gini of the feature *i*. The ratio of the sums in the second fraction measures the frequency with which feature *i* was selected when it was evaluated.

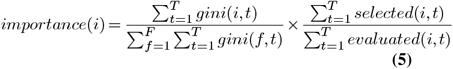

### D. Model evaluation

Standard metrics (accuracy, precision, recall, F1-score) are used to compare the classification performance of RFs and LAVASET. For both the simulated and MIBS cohort inputs, LAVASET and RF were run 20 distinct times (20 run pairs with 100 and 1,000 trees, respectively). Identical random state seeds were assigned per run pair for the sample bootstrapping, to account for the randomness between the comparisons. The data were split into training and test sets (80% and 20% respectively), with the sets always being kept identical across the runs. Mean accuracy, precision, recall and F1-scores are reported for the 20 runs.

We performed a grid search optimising the number of trees and neighbours for the MIBS cohort. Tree values ranged from 100 to 10,000 and neighbours from 1 to 50, and all possible combinations were evaluated. The optimal value for the number of trees for LAVASET was 1,000, this model is referred to as LAVASET-1K. We compare LAVASET-1K to two RF models: the first is an RF with the same number of 1,000 trees, referred to RF-1K hereafter, and the second is an RF that runs for the same amount of computational time as LAVASET. We evaluated the number of RF trees that can be fit in the same amount of time as LAVASET-1K required. LAVASET-1K fits 1,000 trees for 18 neighbours (optimal for MIBS cohort) in approximately 11 minutes (on an HP Z6 G4 workstation with 16-core Intel(R) Xeon(R) Silver 4110 CPU @ 2.10GHz with 128GB RAM), a classic RF can include approximately 41,000 trees in the same amount of time (referred to as the RF-41K model).

Precision and recall scores are also calculated as metrics for evaluating the peak coverage of the feature importance performance. In the simulated dataset, the peak points used to create the two distinct groups are assigned as the ground truth (positive designation) for the peak coverage. To evaluate the specificity of LAVASET, we include points outside the peak (equal to the k neighbours assigned) in the calculations of precision and recall. These serve as the true negative designations. The threshold for positive or negative designation in these calculations is whether the point has been assigned a feature importance value (feature importance*>*0) or not (feature importance=0).

The neighbours for the VCG data were calculated on the basis of the time of the cardiac cycle. I.e. selecting Frank’s lead x for variable (time) i also includes the other 2 leads (y, z) at time i. Including more neighbours takes place in steps of 6 (x, y and z for both i *±* 1). Where the first and last time points are also considered neighbours at i *±* 1.

Finally, we assessed LAVASET’s performance on feature extraction for DR, specifically for the 3D CMR dataset. Due to computational constraints, the 3D dataset was tested on 100 and 200 neighbours and 150,000 trees. The neighbours were decided by the x, y, and z coordinates (spatial distance) in an iterative manner by considering the 100 neighbours (for each point) that remain neighbours for over half of the 1,273 samples. The most important features (above the 50% importance value threshold) were selected for LAVASET and RF models (both 150,000 trees), and for each set, we performed further DR and clustering to evaluate how the clusters separated the HCM and HC samples. Uniform manifold approximation and projection (UMAP) and k-means clustering with k=2 (expected number of groups) were used in both cases (16). UMAP components and k-means transformations were evaluated for 20 different random state restarts to ensure robustness. Each restart produced 2 clusters that were scored by the standard metrics, to determine how closely they matched the true sample classifications.

## Results

### E. Simulated NMR dataset

LAVASET yields a non-inferior prediction accuracy compared with traditional RFs when predicting the two simulated groups in the NMR dataset (accuracy: 0.859 *±* 0.027 LAVASET, 0.823 *±* 0.021 RF, Supplementary Figure 5). As shown in Supplementary Figure 6, the RF method (panels B and D) fails to identify all individual features relating to the same peak(s) and instead only identifies a subset of these. In contrast, LAVASET not only does identify correctly the simulated metabolites, but also assigns appropriate importance values to all or most of the correlated points that encompass a peak. Specifically, for ethanol’s CH_2_ peak shown in Figure 2A there is 92% precision and 100% recall in capturing the peak points. For the CH_3_ peak precision is 77% and recall 95%. In contrast for the RF model, we see much lower recalls (CH_2_ peak: 0.36, CH_3_ peak: 0.18) but 100% precision given that the model simply does not capture almost any points on or around the peak. For uracil, the second multi-peak metabolite used to create the simulated dataset, we see that LAVASET has 100% recall in both doublets and 92% and 80% precision respectively. RF underperforms in capturing all the points of these metabolites as well with recalls at 32% and 47%.

**Fig. 2.**
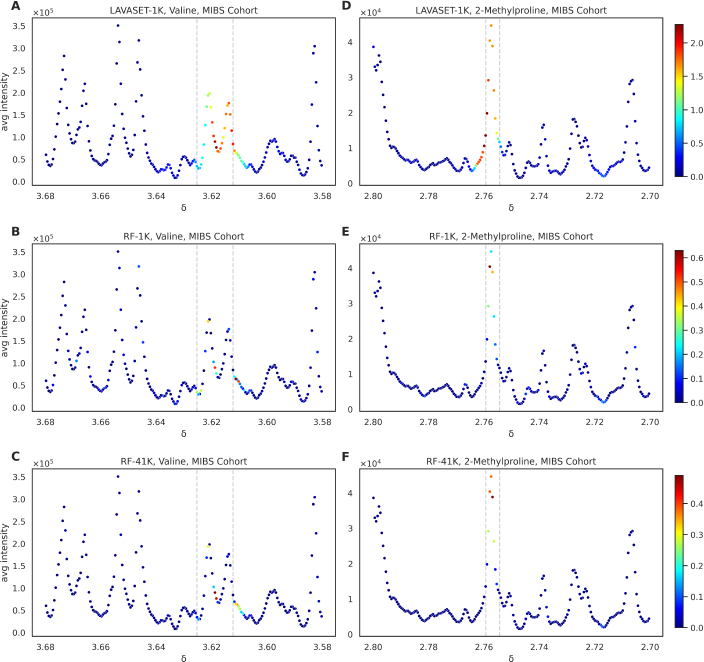
Comparison of LAVASET-1K to RF-1K and RF-41K feature importance assignments on pre-identified metabolites. Panels A, B, C shows feature importances (as defined in Equation 2 in Methods) for valine, and panels D, E, F for 2-methylproline. Colourbar on the right indicates the feature importance value range (red = higher, blue = lower). Dashed gray lines indicate the previously identified points for the specific metabolites peaks. X-axis indicates the chemical shift (in parts per million (ppm), *δ*). Y-axis shows the average signal intensity.

### F. MIBS cohort

After a grid search evaluation to identify the optimal parameters for the number of neighbours and trees, the highest scoring accuracy was achieved by 1,000 trees and 18 neighbours (Supplementary Figure 7). Average accuracy values overall dropped after considering more than 20 neighbours and there were no significant differences when taking more than 1,000 trees for the specific dataset. Supplementary Figure 8A presents the optimal number of neighbours for the 1,000 trees with a clear peak being displayed on the graph, while the drop after 20 neighbours is also evident there. LAVASET yields similar results when classifying IBS vs HC in the MIBS cohort (only for this cohort, losing 1% in mean accuracy in comparison to RF). Across 20 runs the average accuracy score for LAVASET is 0.68 *±* 0.02, for RF-1K 0.69 *±* 0.01, and for RF-41K 0.68 *±* 0.02. Supplementary Figure 8B boxplots represent these accuracy values, along with precision, recall, and F1 score values. The value ranges shown for the RF-41K model suggest that when running 41,000 trees there is potential for over-fitting, given that the values across each of the 20 runs are almost identical. For the 4 different performance metrics the error bars of LAVASET vs RF-1K and LAVASET vs RF-41K overlap, indicating LAVASET is non-inferior to either RF model (Supplementary Figure 8B).

The most pronounced differences between LAVASET and RF are evident when looking at the feature importance. The ability of LAVASET to capture the entirety of peaks attributed to a metabolite surpasses the RF feature evaluation which merely captures less than half of the points that encompass the peak. Figure 2 shows the relevant peaks of previously identified metabolites valine (A, B, C) and 2-methylproline (D, E, F). These two metabolites have been previously associated with separating IBS from HC patients, by showing high feature importance in classification models using Support Vector Classifiers (6). Panels A and D show clearly how LAVASET is able to assign importance values to all points of the valine and 2-methylproline peaks, respectively. The dashed gray lines indicate the previously identified point ranges for the specific metabolites peaks, which present a ground truth for the peak, but can sometimes be affected by small shifts or missing points on the sides of a given peak. In the case of 2-methylproline we notice that LAVASET is also capturing and assigning relatively high importance on points to the left side of the previously-known peak. These points, however, when visualised on the spectrum appear to be part of the rest of the peak and LAVASET only assigns an importance to the points up to where the next peak is starting, without including that next non-related peak. This can also be seen on the right-side of valine in Figure 2A, however in this case importance is relatively lower, presenting a gradient trend as we are reaching the end of the visualised peak.

RF, on the other hand, assigns feature importance to sporadic points of the peak, with no real continuance as to the values of importance (valine, Figure 2B, red point indicating high importance next to light blue point indicating more than half of an importance value). Even in the case of the 41K trees we see that the points selected remain almost the same as in the 1K trees, suggesting that an RF cannot reach LAVASET’s ability in capturing whole peaks, even if the number of trees increases 41-fold. Overall, LAVASET’s output is quite stable and not affected significantly by small changes in the number of neighbours. Peak coverage is equivalent when looking at 10-16 neighbours, with the main differences occurring in the value of importance to each point (Supplementary Figure 9).

### G. MTBLS1 cohort

LAVASET was further tested on the MTBLS1 T2DM cohort, as described in Methods, to ensure that performance stability and feature importance interpretation remain effective in other cohorts. Consistent with previous examples, LAVASET presents non-inferior results to RF (LAVASET accuracy: 0.82 *±* 0.01, RF accuracy: 0.77 *±* 0.02, Supplementary Figure 10) and manages to capture the metabolites pre-identified by (7). Metabolite peak capturing by LAVASET is again superior to RF, by encompassing if not all, most points and attributing feature importance values more equally. Results are expanded in Supplementary Figure 11.

### H. ECG data

We assessed LAVASET’s capabilities on more complex data types where it is common practice to use transformations of the raw reading data to infer further information. An example of such data types is transforming Electrocardiogram (ECG) readings to Vectorcardiogram (VCG) inputs. For this task, we used the ECG data from the Physikalisch-Technische Bundesanstalt (PTB) dataset (8), as described in Methods. LAVASET-1K performed similarly to the RF-1K in this dataset, with accuracy values showing identical results across 20 runs (0.81 *±* 0.01, Supplementary Figure 12) and other metrics having only small differences (Supplementary Figure 12). Figure 3 shows the results when taking 20 nearest neighbours (across the time axis as described in Methods) and 1,000 trees. LAVASET is able to capture peaks and anomalies across the ECGs that indicate differences between the HC and MI cases. Given the nature of the dataset and the idiosyncrasies of different MI cases based on the location of the infraction, setting a binary classification by incorporating all types of MI as one target will only capture important features related to all these different MI phenotypes.

**Fig. 3.**
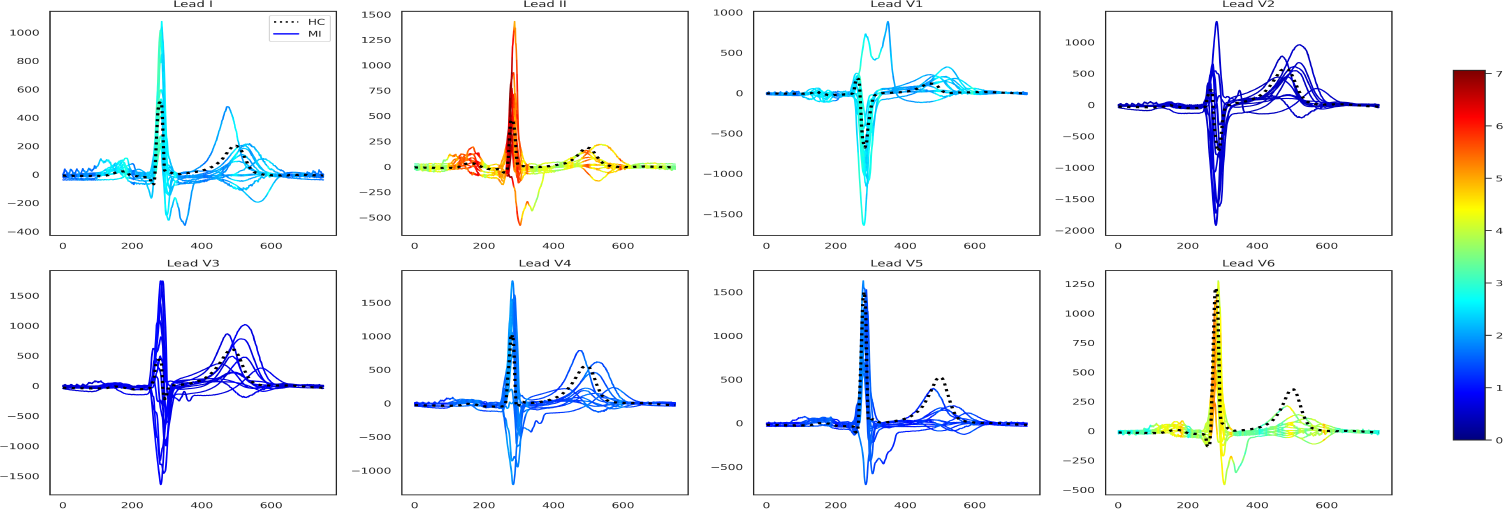
ECG feature importance (Kors regression back-transformed from the VCG feature importance) normalised across the 8 independent ECG leads. Subplot titles indicate the respective leads. X-axes show the time in milliseconds (ms) and y-axes the voltage magnitude in millivolts (mV). The dotted black line indicates a healthy sample and the solid lines represent the 10 MI types (acute, anterior, anteriolateral, anteroseptolateral, anteroseptal, inferior, inferolateral, inferoposterolateral, lateral, posterior) in this dataset and are coloured by relative feature importance.

### I. CMR Imaging

Both the LAVASET and RF models demonstrate equivalent values across all metrics. LAVASET shows an accuracy of 0.878, precision of 0.837, recall of 0.904 and F1-score of 0.869. RF shows an accuracy of 0.875, precision of 0.916, recall of 0.851, and F1-score of 0.882. Given these high values, we are confident about the use of both models for feature extraction, as means to DR. Figure 4 shows the results on 100 neighbours and 150,000 trees. LAVASET is able to assign importance values to a larger surface area of the left ventricle, while also demonstrating the most important areas. RF, on the other hand, picks patches of the left ventricle as most important, disregarding other anatomical parts that could be important. To test how informative these selected features are, we used them as input in further DR and clustering via UMAP and k-means. Table 1 shows the clustering performance across 20 iterations of UMAP and k-means, indicating that the features selected by LAVASET perform constantly better than those selected by RF. We also highlight that in this specific dataset, feature extraction improves the models considerably, given the significantly lower performance observed when taking all features as input.

**Table 1.**
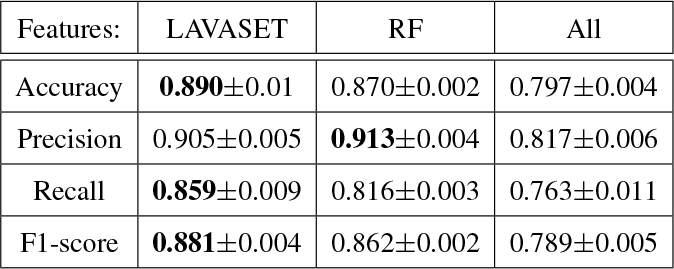
HCM CMR dataset, feature DR performance. Scores for accuracy, precision, recall, and F1-score across 20 iterations of UMAP and k-means (k=2) on the selected feature sets by importance in LAVASET, RF and without selection. Values shown are the mean*±*standard deviation.

**Fig. 4.**
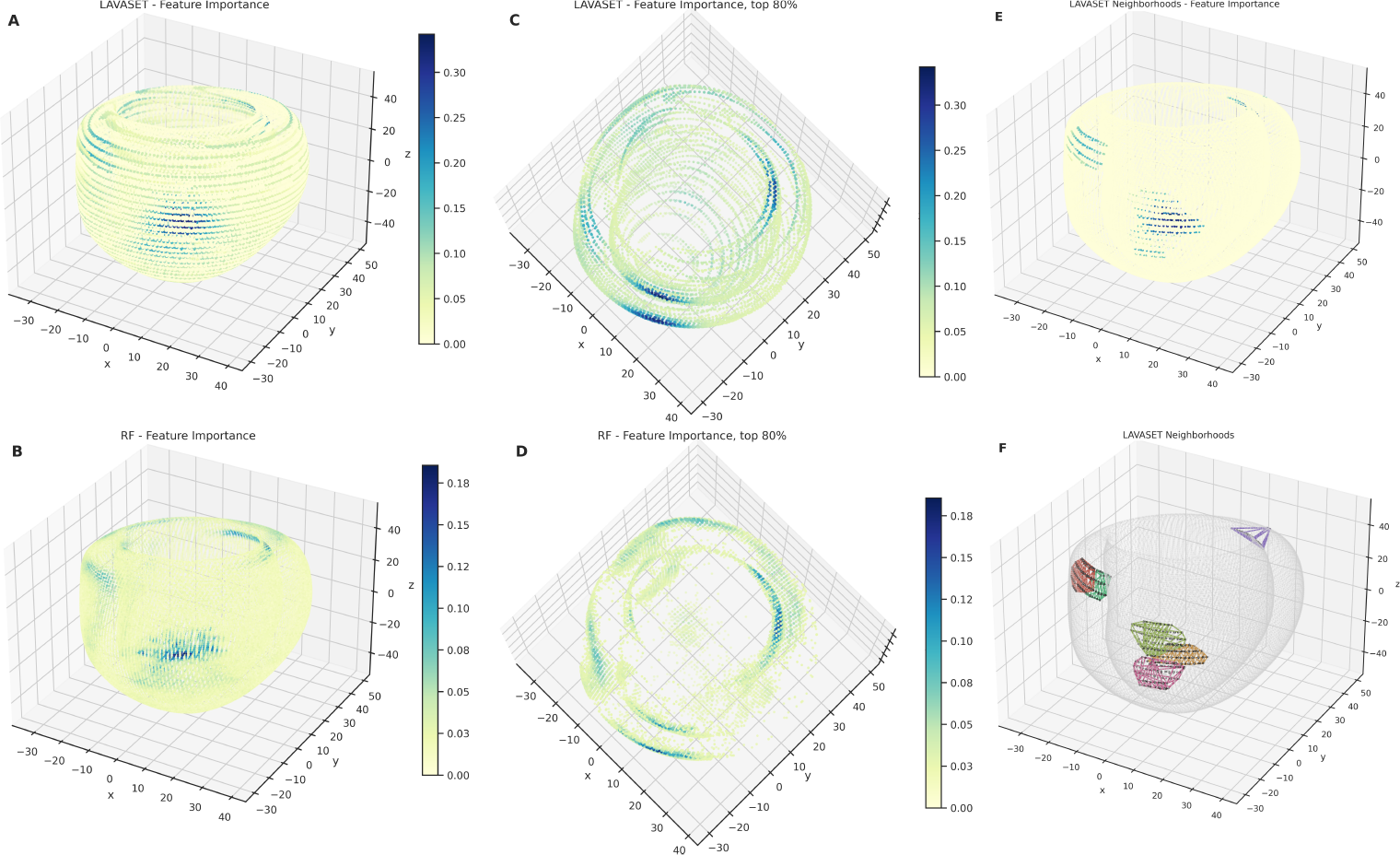
3D representations of the left ventricle CMR data points (x, y, z coordinates). Points represent an averaged template of HCM and HC ventricles. The inner and outer structures formed show the endocardium and epicardium, respectively. Colourbar shows the feature importance gradient, indicating that in panel A (LAVASET) the assignment of higher importance is encompassing the entirety of the ventricle structure. In panel B, assignments are given in a patch-like manner for RF. Panels C and D show the points in the 80% quantile from a top view to facilitate the distinction between the inner and outer walls of the ventricle. Panels E and F show 6 distinct neighbourhoods of FOIs. In panel D convex hulls are drawn for each neighbourhood to represent the pattern of points per neighbourhood. Black points indicate the neighbours and the coloured connecting lines emphasise the different neighbourhoods, and the string-like pattern of neighbour points. Panel C shows the feature importance for the respective 6 neighbourhoods.

## Discussion

We have presented a novel ensemble learning method that aims to enhance variable importance and ensure that correlated features are evaluated appropriately and not independently. LAVASET produces non-inferior performance results to traditional RFs in all but one of the examples tested above, and in both simulated and real datasets. While not sacrificing performance in most examples, it is able to assign feature importances in a superior way, by not missing features that are as ‘important’ by means of correlation, and ensuring that importance values are correctly distributed. In contrast, we have shown that RF allocates feature importance to isolated points, whether that is peak points in a spectrum or points on a 3D mesh, without any clear consistency of the magnitude of importance. Even when substantially increasing the number of trees in RF, the selection of points and assignment of importance are nearly identical to those selected with the least number of trees, and still do not capture the points selected by LAVASET.

The motivation behind developing LAVASET stems from the idea of enhancing traditional RFs, in a manner similar to how Group Lasso enhances the Lasso algorithm. In the main premise of Group Lasso, we also assume that there are groups of features that are expected to have similar effects or are related to each other. By penalising the sum of the absolute values (L1 norm) of the coefficients within each group, Group Lasso encourages the model to select entire groups of features together or exclude them altogether (17). Like in Group Lasso, LAVASET is particularly useful when dealing with high-dimensional data where groups of features exhibit similar importance or are structured in some meaningful way. This was evident by the variety of datasets we tested, where the number of features ranged from 18,000 to 46,000. In this high-dimensional context, LAVASET exhibits stability in its output and remains relatively unaffected by minor variations in the number of neighbours comprising the groups. However, Group Lasso requires non-overlapped groups of features, whereas in LAVASET this can be varied. In fact, in LAVASET, different FOIs can have different numbers of neighbours to increase flexibility. Likewise, LAVASET emulates the kernel filter in convolutional neural networks (CNNs) in that it combines multiple features into a single output (for splitting in LAVASET), however, it does so without condensing the output and attributing the feature importance across the initial features. Other work has investigated the relations between individual features in terms of the similarity of performance at different splits. This methodology is able to discern correlations between individual features, however the mutual forest impact is constrained to evaluating pairs of features only (18). LAVASET can, in theory, be combined with this to perform an a posteriori analysis of evaluating correlations between groups of latent variables. LAVASET’s main limitation derives from the requirement of defining the aforementioned ‘groups’ (akin to the kernel size in CNNs). This parameter is user-specified and assumes an understanding of the relationship between the variables. In the examples we presented here, this relationship is defined and assigned by distance, whether that is 1D distance across a spectrum, 1D across time-series data, or 3D spatial distance. The influence of the neighbourhood definition is evident, especially in the 3D CMR example. Figure 4 clearly shows that the string-like pattern of importances calculated by LAVASET is predominantly driven by the assignment of neighbours for each FOI (Figure 4F). This inherent limitation, however, is what enables LAVASET’s flexibility in creating the groups of neighbours. Distance is only one of the metrics that can be employed. A few other potential examples include genomic distance (combine SNPs via linkage disequilibrium), mass spectrometry isotope patterns (proteomics, metabolomics), or hierarchical relationships (e.g. taxonomy of microbiota). This attribute renders LAVASET more versatile than other similar methods, while giving the user the ability to tailor the algorithm to their specific needs. Hence, it is applicable to a wide variety of datasets and biological questions.

This flexibility is also translated to LAVASET’s code implementation. The algorithm also exploits parallelisation in order to speed up computations. The body of the code is written in Python 3.10, while utilizing established C++ scripts for efficiency. Given the nature of the code and algorithm, LAVASET can be run in batches, if needed, or can be easily altered to incorporate additional metrics to distance.

To enhance and expand LAVASET’s capabilities, we are working on incorporating the gradient boosting algorithm as one of our built-in additional components. This will extend LAVASET’s core methodology to other ensemble methods and benefit from iterative learning and the specific advantages of boosting trees.

## Conclusion

We have presented LAVASET, a novel ensemble method capable to improve feature interpretability by selecting relevant groups of features instead of individual features. Its novel functionality is most useful in datasets with correlations between features. In cases where this is missing from the data, then traditional RFs are more appropriate. LAVASET offers interoperability to the user both by the structure of its code and via the neighbours parameter. It can be applied to almost all omics data types to identify all relevant known or unknown important features and effectively perform feature extraction for DR.

## Acknowledgements

Conceptualisation: JMP and MK. Methodology: JMP, MK and KX. Investigation: MK, JMP and KX. Visualisation: MK and JMP. Writing - original draft: MK and JMP. Writing - editing: MK, JMP, TE, KX, JSW and DPO’R. Supervision: JMP, TE and JSW. We thank Zlatan Mujagic (6) for providing the raw MIBS cohort NMR data, the DPO’R group for supplying the CMR images, and Sanjay Prasad for the RBH CMR data.

## Supplementary

**Fig. 5.**
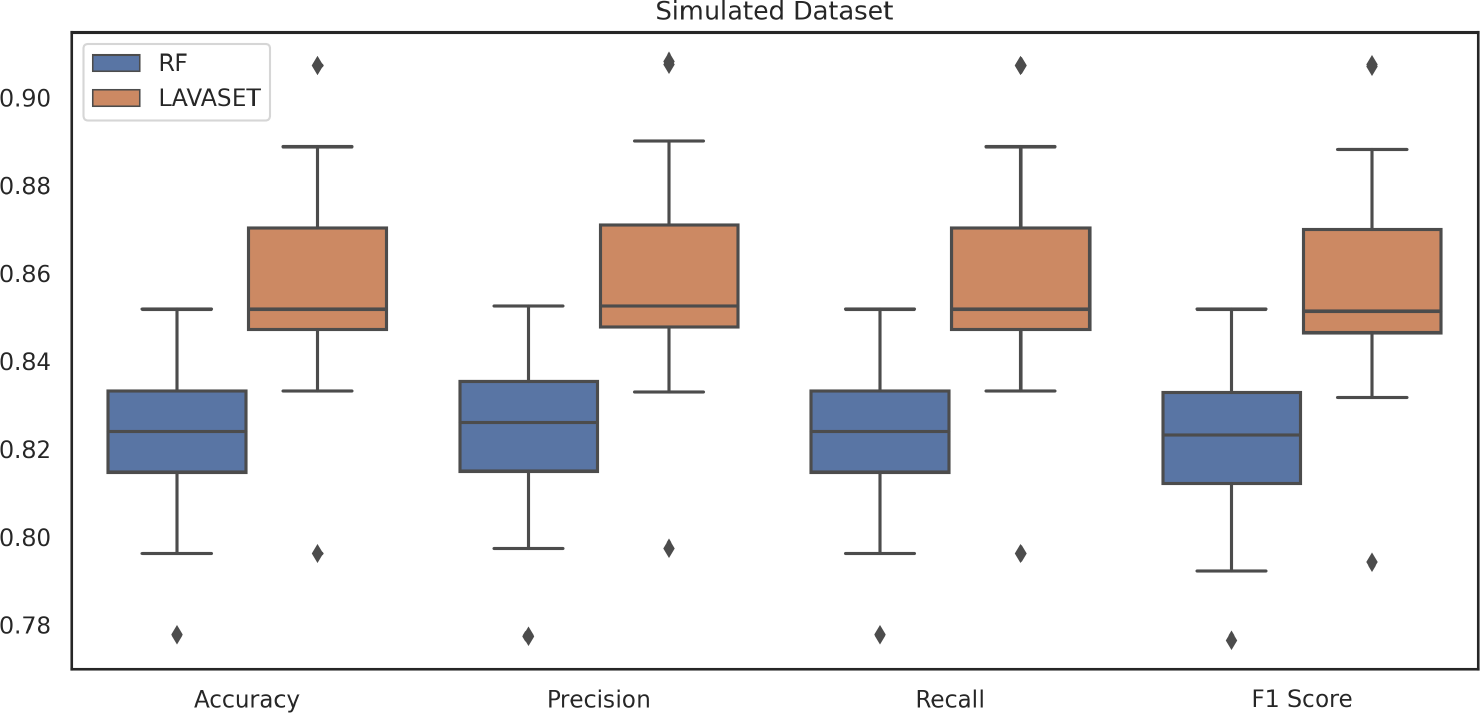
Performance in the simulated dataset. Models were run for binary classification on simulated groups generated from the MIBS dataset. LAVASET outperforms RF across all metrics as seen in 20 distinct iterations with identical random seeds between LAVASET and RF. Accuracy: LAVASET: 0.859*±*0.027, RF: 0.823*±*0.021. Precision: LAVASET: 0.860*±*0.026, F: 0.824*±*0.020, Recall: LAVASET: 0.859*±*0.027, RF: 0.823*±*0.021, F1 Score: LAVASET: 0.859*±*0.027, RF: 0.823*±*0.021.

**Fig. 6.**
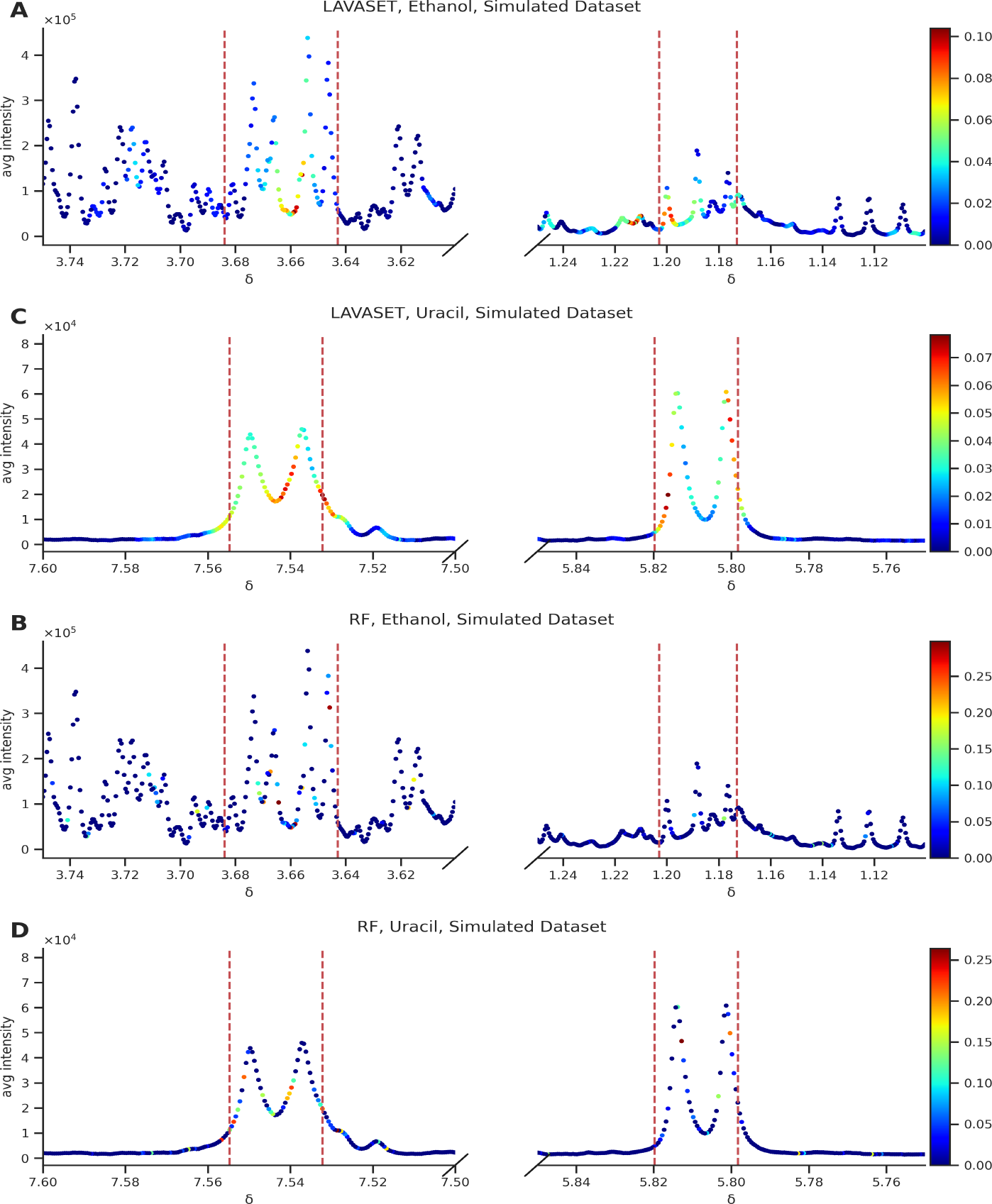
LAVASET feature importance assignments on the simulated dataset. A close-up look at the specific spectral widths for ethanol and uracil, the top metabolites used to separate two groups in this simulated dataset. LAVASET shows better performance in capturing most points of the peak versus the RF method which identifies less than half. Specifically: Ethanol - LAVASET: CH_2_ peak: precision: 0.92, recall: 1.00, CH_3_ peak: precision 0.77, recall: 0.95, RF: CH_2_ peak: precision: 1.00, recall: 0.36, CH_3_ peak: precision: 1.00, recall: 0.18, Uracil - LAVASET: H-6 (next to C=O) doublet: precision: 0.92, recall: 1.00, H-5 (next to NH) doublet: precision: 0.80, recall: 1.00. RF: H-6 doublet: precision: 1.00, recall: 0.32, H-5 doublet: precision: 0.94, recall: 0.47. Colourmap bar to the right indicates feature importance Gini values, x-axis is marked by the parts per million metric (ppm or *δ*). The simulated data were run on 100 trees.

**Fig. 7.**
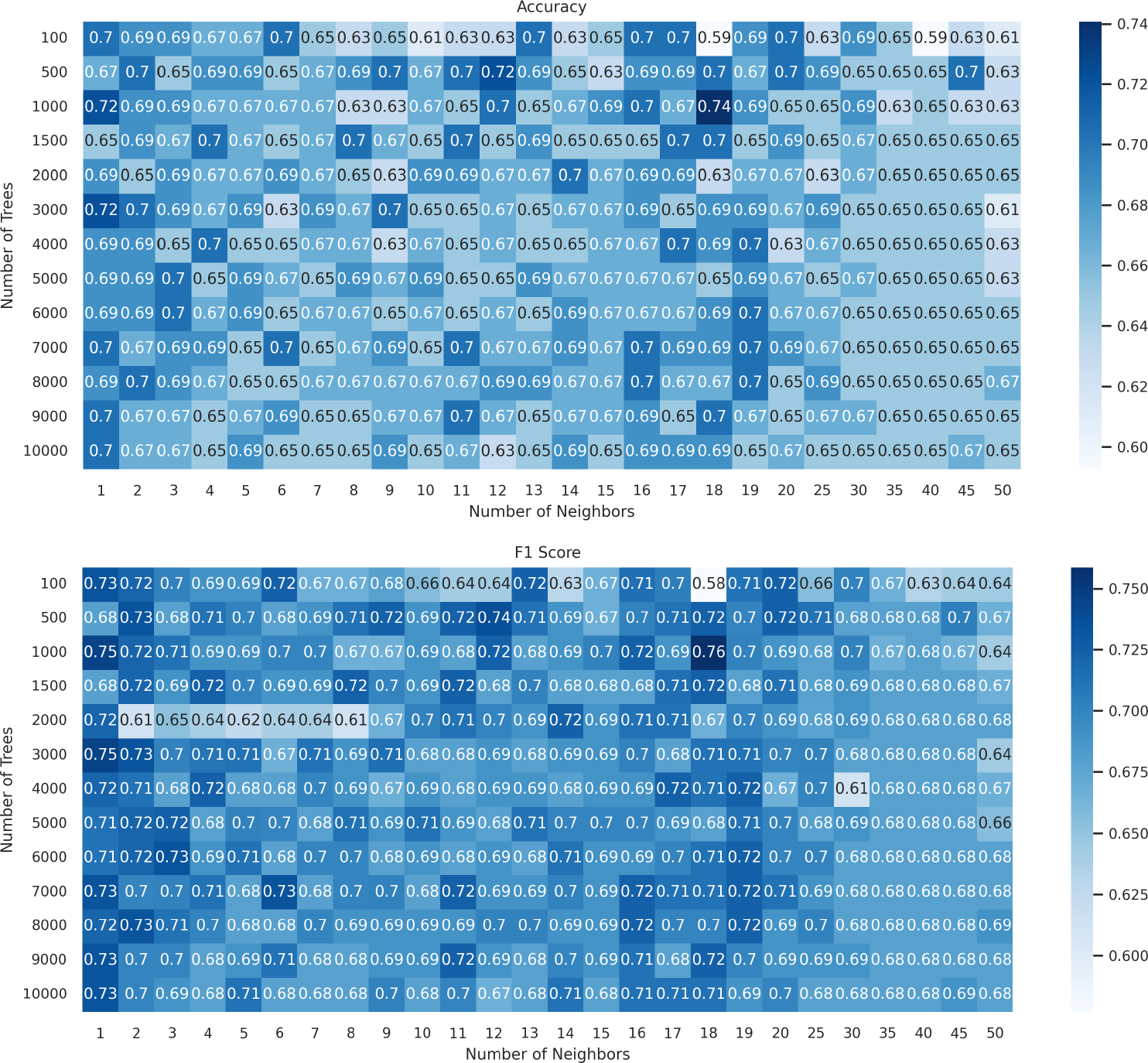
Accuracy, precision, recall and F1-score values for grid search parameter tuning in MIBS dataset, when performing binary classification of IBS vs HC.

**Fig. 8.**
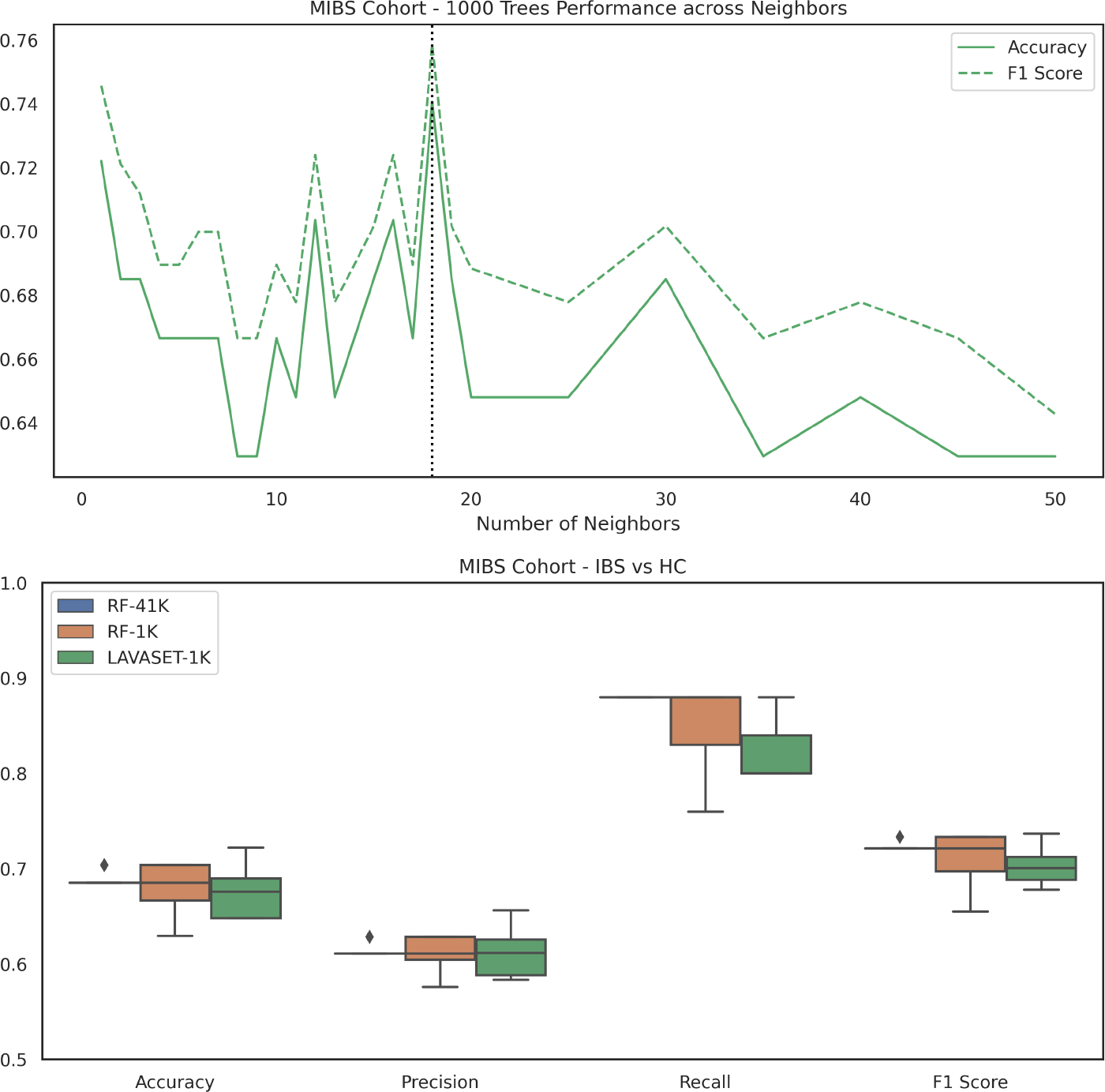
Panel A shows the changes in performance when looking at a different number of neighbours, evaluated by LAVASET-1K trees, in the MIBS cohort. Solid line presents the accuracy and dashed line presents the f1 score, the black vertical line indicated the best-performing number of neighbours at k=18. Panel B compares the performance of the LAVASET 1,000 trees / 18 neighbours (green) to the RF-1K trees model (blue) and the RF-41K model (time comparison, orange).

**Fig. 9.**
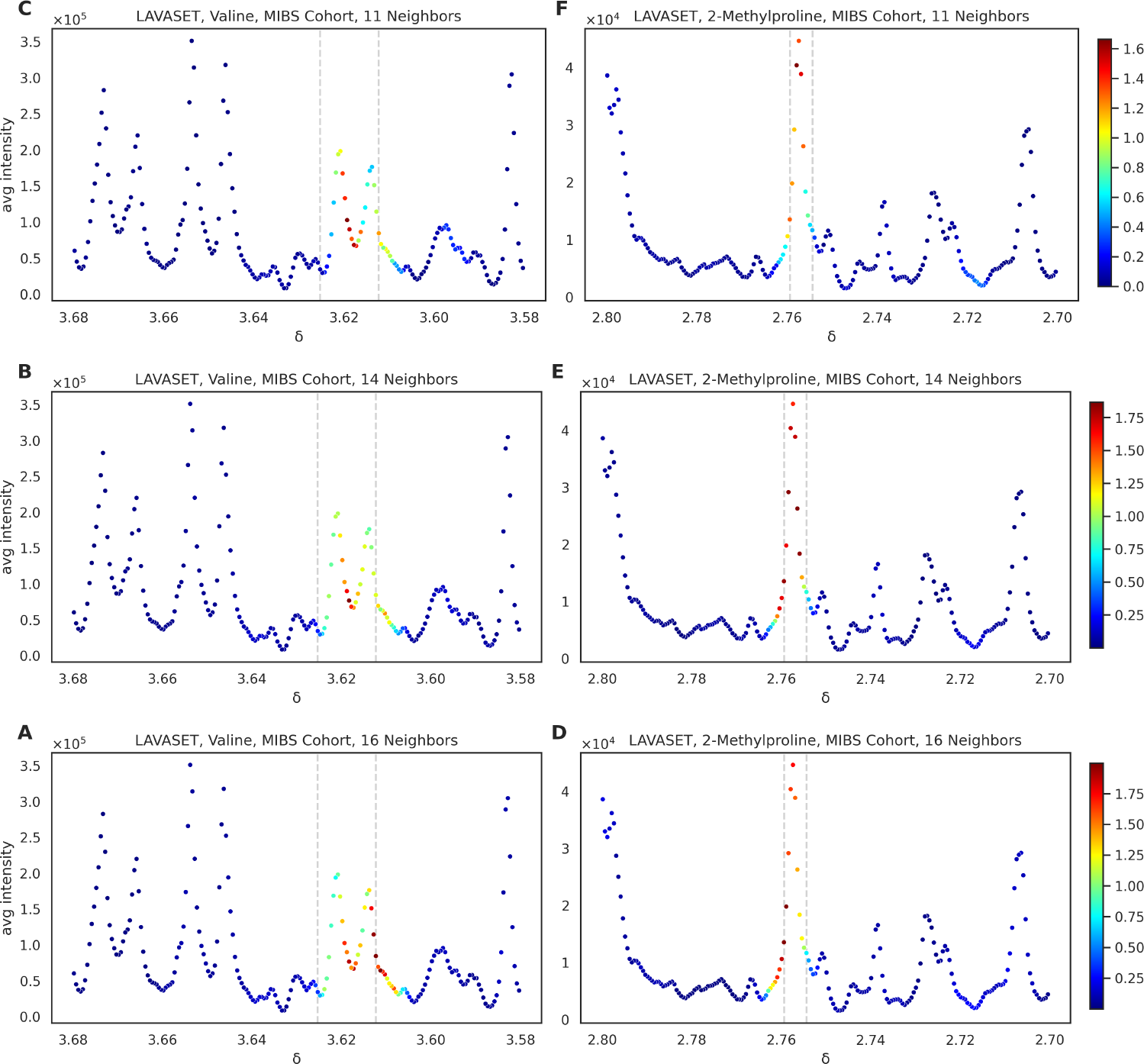
Feature importance coverage across different number of neighbours for the MIBS cohort.

**Fig. 10.**
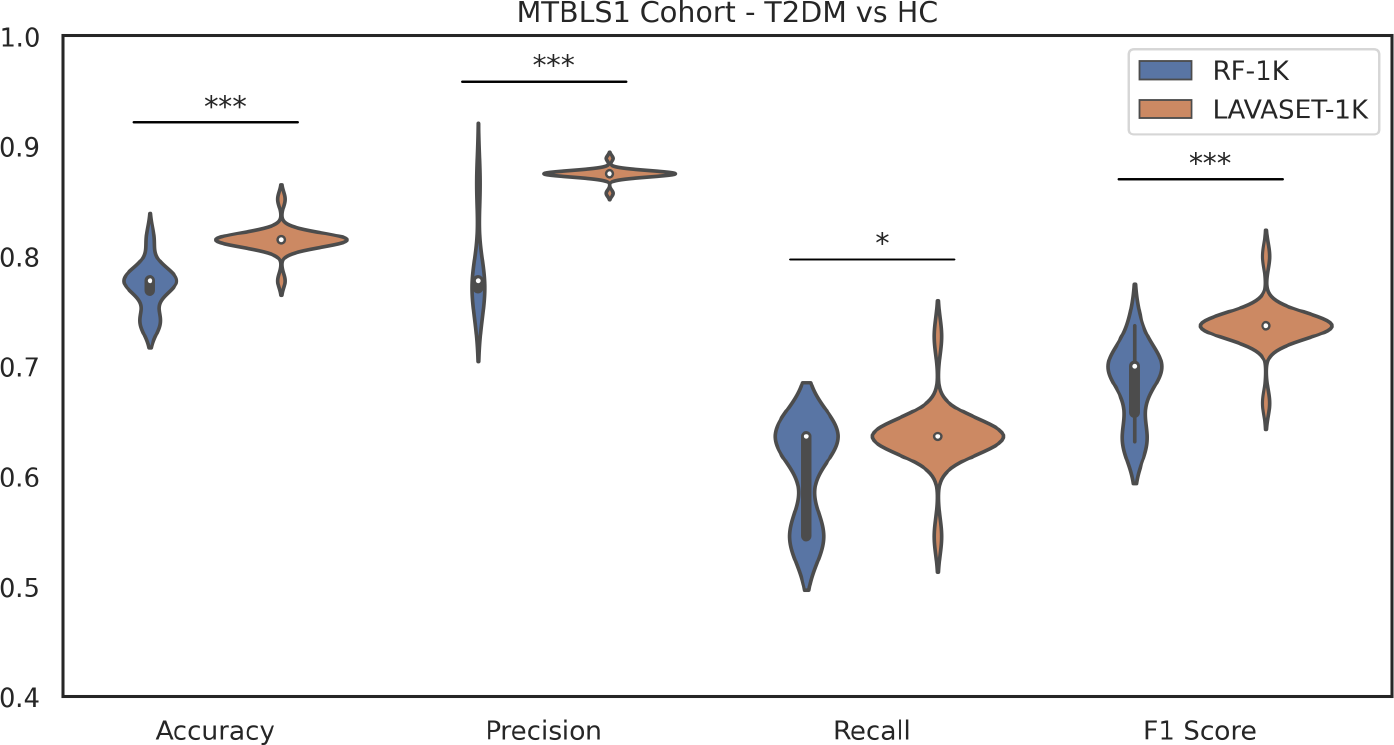
Performance in the MTBLS1 dataset. Models were run for binary classification between Type 2 Diabetes Mellitus (T2DM) samples and Healthy Controls (HC). LAVASET outperforms RF across all metrics, as seen in 20 distinct iterations with identical random seeds between LAVASET and RF. Accuracy: LAVASET: 0.82*±*0.01, RF: 0.77*±*0.02, Precision: LAVASET: 0.88*±*0.01, RF: 0.79*±*0.04, Recall: LAVASET: 0.64*±*0.03, RF: 0.61*±*0.04, F1-score: LAVASET: 0.74*±*0.02, RF: 0.68*±*0.03.

**Fig. 11.**
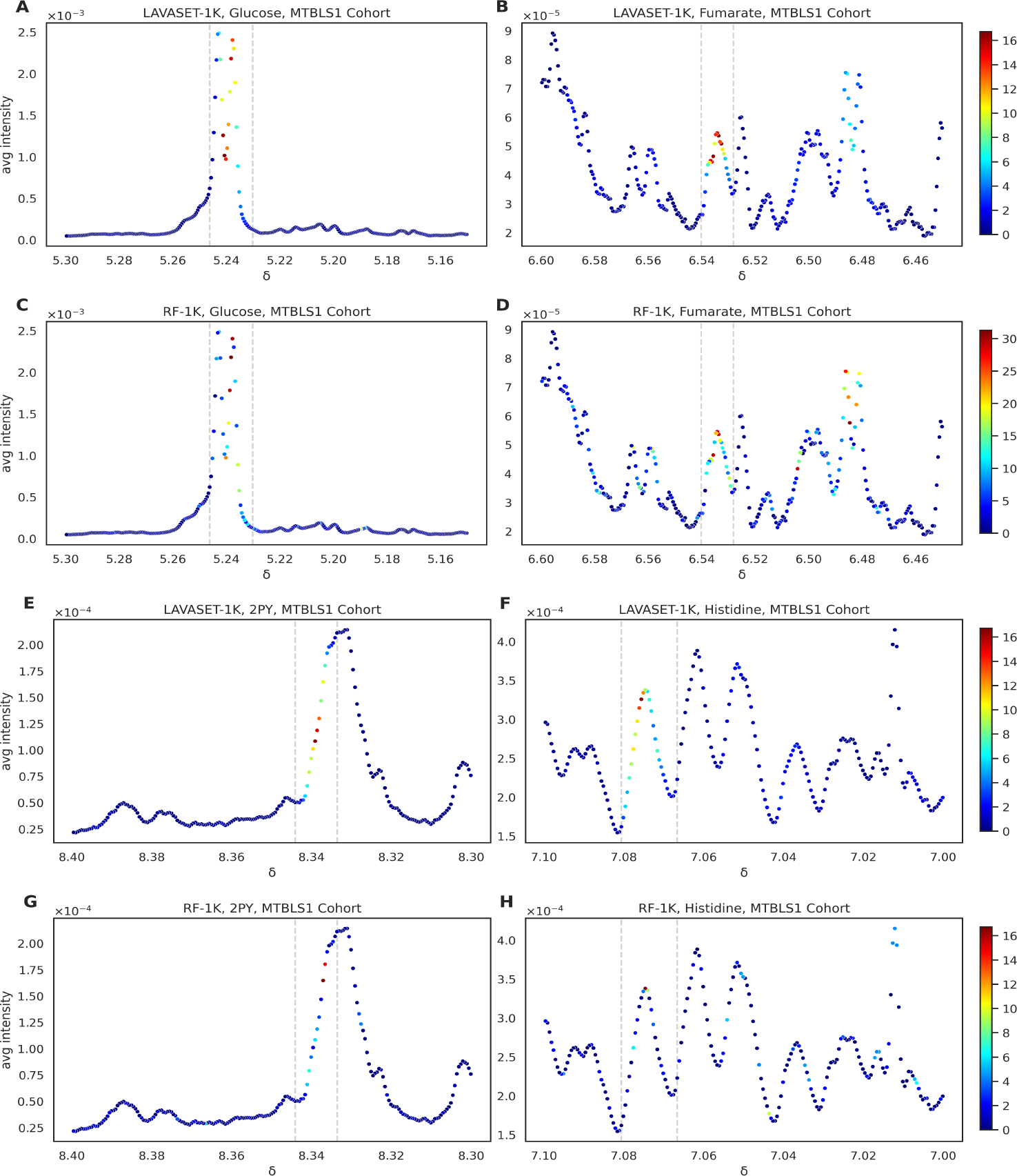
Feature importance coverage in the Type 2 Diabetes Mellitus (T2DM) MTBLS1 dataset. All figures show results of LAVASET and RF models across 1,000 runs and 10 neighbours (for LAVASET). Panels A and C present glucose coverage in LAVASET and RF respectively, as a peak capturing baseline given the classification was done between T2DM samples and HC. Panels B/D present fumarate, E/G *N*-methyl-2-pyridone-5-carboxamide (2PY) and F/H histidine, all three metabolites are identified in the original cohort study. Gray dashed lines represent the ppm values reported in the study. Additional points around the peak are shown to capture LAVASET’s ability of not assigning high importance values on points around the peak (noise). This is evident in the fumarate and histidine examples where RF captures points outside the identified peak ppm values (D, H), while LAVASET evidently assigns much lower importance to those noise features (B, F).

**Fig. 12.**
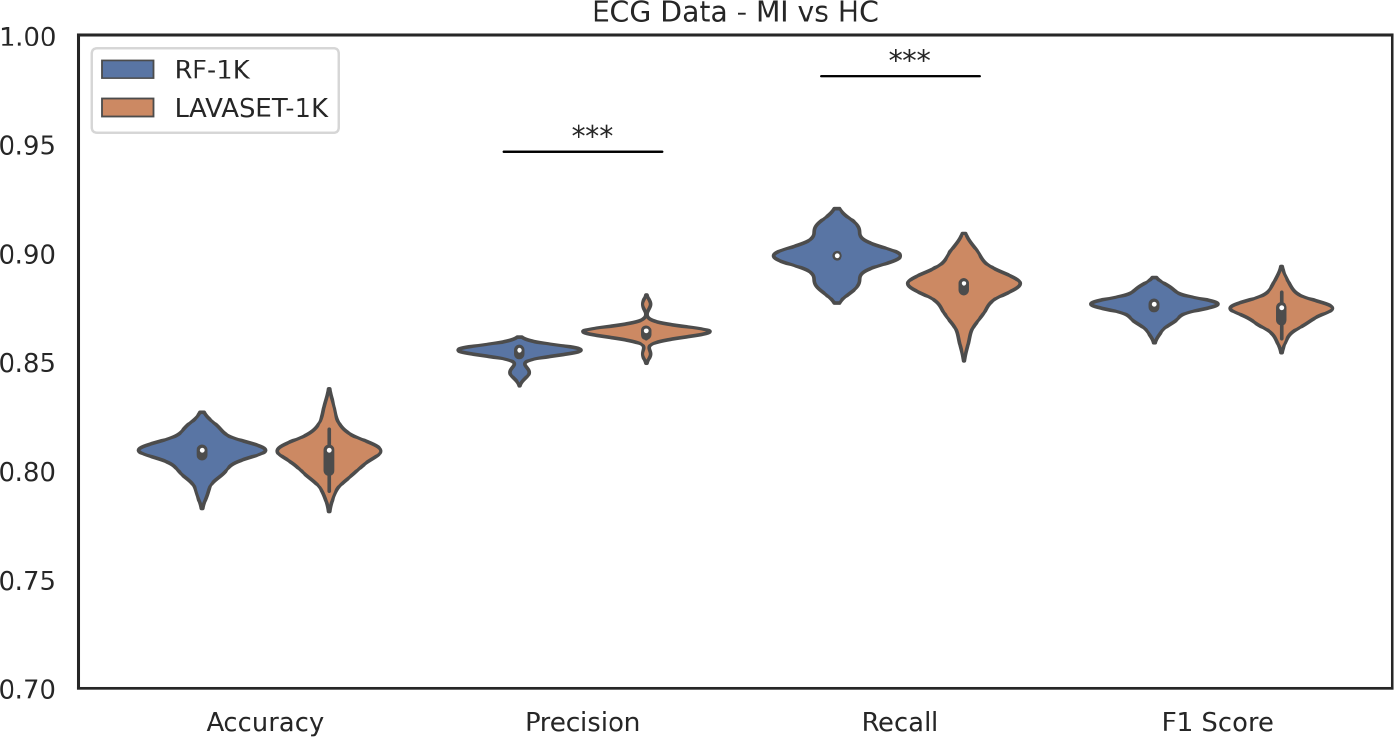
Performance in the ECG dataset. Models were run for binary classification between Myocardial Infraction (MI) samples and Healthy Controls (HC). Accuracy: LAVASET: 0.81*±*0.01, RF: 0.81*±*0.01, Precision: LAVASET: 0.86*±*0.01, RF: 0.85*±*0.01, Recall: LAVASET: 0.88*±*0.01, RF: 0.90*±*0.01, F1-score: LAVASET: 0.87*±*0.01, RF: 0.88*±*0.01.

**Fig. 13.**
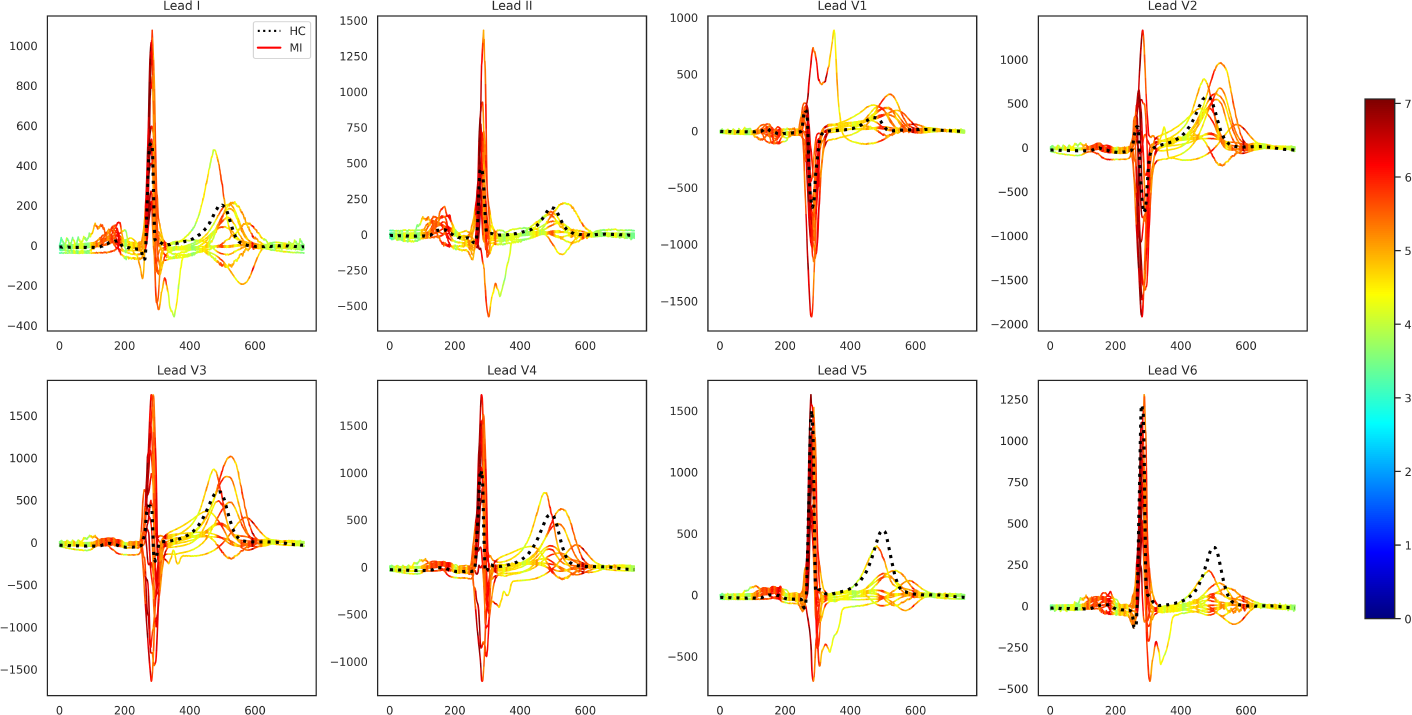
Feature importance coverage in the ECG dataset. Values are normalised per lead, in order to show the most important areas in the individual leads. Black, dotted lines indicate the (average) Healthy Controls (HCs) and green-yellow-red heatmap-coloured lines show all Myocardial Infarction (MI) cases.

